# Co-expression networks reveal the tissue-specific regulation of transcription and splicing

**DOI:** 10.1101/078741

**Authors:** Ashis Saha, Yungil Kim, Ariel D. H. Gewirtz, Brian Jo, Chuan Gao, Ian C. McDowell, GTEx Consortium, Barbara E. Engelhardt, Alexis Battle

**Author notes:** Denotes equal contribution (Yungil Kim, Ariel D. H. Gewirtz).

## Abstract

Gene co-expression networks capture biologically important patterns in gene expression data, enabling functional analyses of genes, discovery of biomarkers, and interpretation of regulatory genetic variants. Most network analyses to date have been limited to assessing correlation between total gene expression levels in a single or small sets of tissues. Here, we have reconstructed networks that capture a much more complete set of regulatory relationships, specifically including regulation of relative isoform abundance and splicing, and tissue-specific connections unique to each of a diverse set of tissues. Using the Genotype-Tissue Expression (GTEx) project v6 RNA-sequencing data across 44 tissues in 449 individuals, we evaluated shared and tissue-specific network relationships. First, we developed a framework called Transcriptome Wide Networks (TWNs) for combining total expression and relative isoform levels into a single sparse network, capturing the complex interplay between the regulation of splicing and transcription. We built TWNs for sixteen tissues, and found that hubs with isoform node neighbors in these networks were strongly enriched for splicing and RNA binding genes, demonstrating their utility in unraveling regulation of splicing in the human transcriptome, and providing a set of candidate shared and tissue-specific regulatory hub genes. Next, we used a Bayesian biclustering model that identifies network edges between genes with co-expression in a single tissue to reconstruct tissue-specific networks (TSNs) for 27 distinct GTEx tissues and for four subsets of related tissues. Using both TWNs and TSNs, we characterized gene co-expression patterns shared across tissues. Finally, we found genetic variants associated with multiple neighboring nodes in our networks, supporting the estimated network structures and identifying 33 genetic variants with distant regulatory impact on transcription and splicing. Our networks provide an improved understanding of the complex relationships between genes in the human transcriptome, including tissue-specificity of gene co-expression, regulation of splicing, and the coordinated impact of genetic variation on transcription.

## Introduction

Gene co-expression networks are an essential framework for elucidating gene function and interactions, identifying sets of genes that respond in a coordinated way to environmental and disease conditions, and identifying regulatory relationships and potential drug targets (Penrod et al., 2011; Xiao et al., 2014; Yang et al., 2014). Each edge in a co-expression network represents a correlation between two transcriptional products, represented as nodes (Stuart et al., 2003). The majority of gene co-expression networks, traditionally estimated from microarray data, have focused on correlation between total gene expression levels, with edges representative of transcriptional co-regulation. However, post-transcriptional modifications, including alternative splicing, have significant effects on the abundance of specific RNA isoforms (Matlin et al., 2005); Mutations that lead to disruption of splicing play an important role in tissue- and disease-specific pathways (Li et al., 2016b; Lee et al., 2012; Wang et al., 2008; DeBoever et al., 2015; Ward and Cooper, 2010; L´opez-Bigas et al., 2005), including tumor progression (Ghigna et al., 2008) and risk of Alzheimer’s disease (Hutton et al., 1998; Glatz et al., 2006). While a number of splicing factors are known, regulation of splicing and the specific regulatory genes involved remain poorly understood relative to the regulation of transcription (Mel´e et al., 2015; Scotti and Swanson, 2015). RNA-sequencing now allows quantification of isoform-level expression, providing an opportunity to study regulation of splicing through network analysis. However, current research estimating RNA isoform-level networks (Li et al., 2014, 2015, 2016a) has not used detailed network representations that distinguish identification of splicing regulation—using relative quantities such as isoform ratios—from transcription regulation—using absolute quantities such as total expression level of each isoform. Moreover, these methods, and initial work on clustering relative quantities, have not be applied to large RNA-seq studies for well-powered network reconstruction in diverse contexts (Dai et al., 2012; Iancu et al., 2015).

Another important gap in our interpretation of regulatory effects in complex traits is a global characterization of gene co-expression networks that are only identified in a disease-relevant tissue type. Per-tissue networks have been estimated for multiple tissues (Pierson et al., 2015; Piro et al., 2011), but have been based on limited sample size. Recent studies have recognized the essential role that tissue-specific pathways play in disease etiology (Greene et al., 2015), but have developed these per-tissue networks by aggregating single tissue samples across multiple studies. However, differences in study design, technical effects, and tissue-specific expression will make the cross-study results difficult to interpret mechanistically, with large groups of genes expressed in similar tissues and studies tending to be highly connected rather than providing detailed network structure (Lee et al., 2004). Most importantly, these analyses do not directly separate effects unique to each tissue from shared, cross-tissue effects.

In this work, we reconstructed co-expression networks from the Genotype Tissue Expression (GTEx) v6 RNA-sequencing data (GTEx Consortium, 2015). These data include 449 human donors with genotype information and 7,051 RNA-sequencing samples across 44 tissues. Here, we identified networks that reveal novel relationships from previous analyses and address two important goals in regulatory biology: identification of edges reflecting regulation of splicing, and discovery of edges arising from gene relationships unique to specific tissues.

To do this, we first built co-expression networks for sixteen tissues, with our framework of Transcriptome-Wide Networks (TWNs), including as variables both total gene expression levels and transcript isoform ratios. Hub genes in these networks with many isoform ratio neighbors were significantly enriched for RNA binding genes and genes known to be involved in splicing, demonstrating the utility of this framework in characterizing regulation of splicing. From the GTEx tissue TWNs, we identified novel candidates of both shared and per-tissue transcriptional regulators of relative isoform abundance and splicing. Next, we built tissue-specific gene co-expression networks across 27 tissues, where each network edge corresponds to correlation between genes that is uniquely found in a single tissue. We show that the TSN nodes with greatest connectivity correspond to genes that are essential for tissue-specific processes. Finally, we confirmed specific network edges from both network analyses by testing associations between a regulatory genetic variant local to one gene with the neighbors of that gene in the co-expression networks. Through this analysis, we provided a list of tissue-specific trans-expression QTLs (trans-eQTLs) and trans-splicing QTLs (trans-sQTLs). Interpretation of regulatory and disease studies will benefit greatly from these networks, providing a much more comprehensive description of regulatory processing including alternative splicing across diverse tissues.

## Results

### Reconstructing Transcriptome-Wide Networks (TWNs) across human tissues

First, we aimed to identify networks that capture a global view of regulation across the transcriptome of diverse human tissues using the GTEx project v6 data. We developed an approach for learning *Transcriptome-Wide Networks* (TWNs) from RNA-seq data, which capture diverse regulatory relationships beyond co-expression, including co-regulation of alternative splicing across multiple genes, and interactions between transcription and splicing. To build a TWN, we first quantified both total expression levels and isoform expression levels of each gene in each RNA-seq sample, and then computed *isoform ratios* (Fig. 1A), representing the relative, rather than total, abundance of each isoform with respect to the total expression of the gene. All values were projected to quantiles of a standard normal distribution. We included both isoform ratios and total expression levels as network nodes, as opposed to estimating a network across all isoform expression levels. This allowed us to distinguish relationships driven by transcriptional regulation from relationships driven by regulation of relative isoform abundance (including alternative splicing). For example, a network over isoform *expression levels* instead of *ratios* would represent the effects of a transcription factor on its target with edges to *all* isoform levels of the target gene, and the same would be true for a splicing factor. In contrast, in a TWN, a transcription factor would only be connected to the total expression node of its target, and a splicing factor to isoform ratio nodes. (Fig. 1C). TWNs can therefore capture, in interpretable form, relationships such as the total expression of a splicing factor affecting relative isoform abundance of other genes (Sveen et al., 2015).

**Figure 1.**
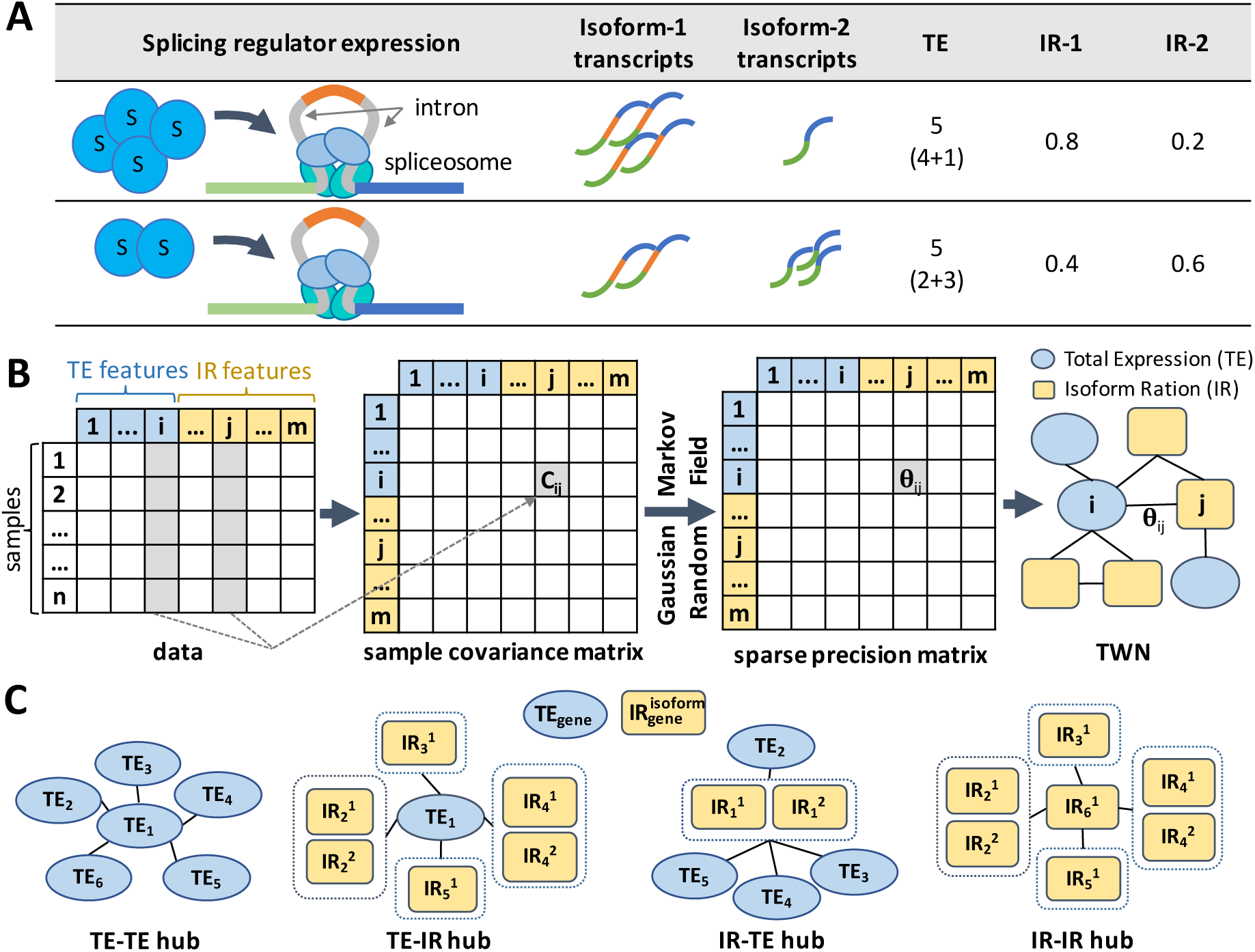
Transcriptome-Wide Network conceptual framework. (A) Schematic of the effect of a splicing regulator on inclusion of a cassette exon, and resulting total expression and isoform ratios of the target gene. Splicing factor expression levels can affect splicing of target genes (Sveen et al., 2015). Higher expression of a splicing regulator S (first row) results in relatively more transcripts of isoform-1 and fewer of isoform-2. Total expression level is constant (5), but isoform ratios are different (0.4 and 0.6) as splicing factor S levels change. (B) The (*i*,*j*)th cell of the sample covariance matrix contains covariance (*C*_*ij*_) between *i*th and *j*th feature in data. We created a sparse precision matrix Θ (inverse covariance) from the sample covariance matrix using graphical lasso to estimate parameters of a Gaussian Markov Random Field. A non-zero value (Θ_*ij*_) in the precision matrix denotes an edge between *i*th feature and *j*th feature in the network. (C) Edges in a TWN represent diverse relationships between total expression (TE) and isoform ratio (IR) nodes. Dotted rectangles group together isoform ratios for different isoforms of the same gene. Of particular focus are network “hub” nodes; in a TWN, there are four possible hub configurations depending on the node type of the central and neighboring nodes.

We then applied graphical lasso (Friedman et al., 2008) to estimate edge weights of a sparse Gaussian Markov Random Field (GMRF) (Rue and Held, 2005) over all nodes jointly, including both the total expression of each gene and the isoform ratio for each isoform (Fig. 1B), Methods). A sparse GMRF captures direct relationships between nodes – a nonzero entry in the precision matrix (interpreted as an edge between two nodes) indicates that the nodes are directly correlated even controlling for effects of all other nodes in the network (i.e., a *partial correlation*) (Sch¨afer and Strimmer, 2005). We modified the basic graphical lasso approach to penalize edges between different node types with different weights, and selected the *ℓ*_1_ regularization parameters for edges between different node types each based separately on the scale-free network property (Methods, Supplementary Table 1, Supplementary Fig. 1-2).

We reconstructed TWNs independently for each of 16 human tissues from the GTEx project v6 data, restricting to tissues with samples from at least 200 donors. We focused on a subset of 6, 000 expression level and 9, 000 isoform ratio nodes for each tissue, based on expression levels, gene mappability, and isoform variability (Methods). We excluded chromosome Y, non-coding genes, and mitochondrial genes. Both technical and biological confounding factors such as batch, RNA Integrity Number (RIN), and sex may introduce correlations among genes in transcription studies (Leek et al., 2010), resulting in numerous false positives in co-expression network analysis (Buettner et al., 2015). Therefore, before applying graphical lasso, we corrected expression data from each tissue for known and unobserved confounding factors using HCP (Mostafavi et al., 2013) (Methods). Additionally, we excluded edges that were unlikely to represent meaningful biological relationships from downstream analysis, including edges connecting gene pairs with overlapping positions in the genome, edges connecting gene pairs with that have been shown to have RNA-seq read cross-mapping potential, and edges between distinct features of the same gene (Methods).

On average, each TWN contained 60, 697 total edges, with 24, 527 strictly between total expression nodes, 18, 539 strictly between isoform ratio nodes, and 17, 631 connecting total expression to isoform ratio nodes (Fig. 2A). We found many “hub” nodes, or nodes with large numbers of neighbors, as expected in biological networks and scale free networks more generally (Barabasi and Oltvai, 2004). Based on a threshold of ten or more neighbors, TWNs had a mean of 1853 “TE-TE” hub genes (total expression nodes connected to many total expression neighbors) and 325 “TE-IR” hub genes (total expression nodes connected to many isoform ratio neighbors) across tissues (Fig. 2A). Hubs with numerous total expression neighbors were more common, but hubs with isoform ratio neighbors were also found in every tissue (Fig. 2A). Complete TWNs for all 16 tissues are available at http://gtexportal.org (in progress).

**Figure 2.**
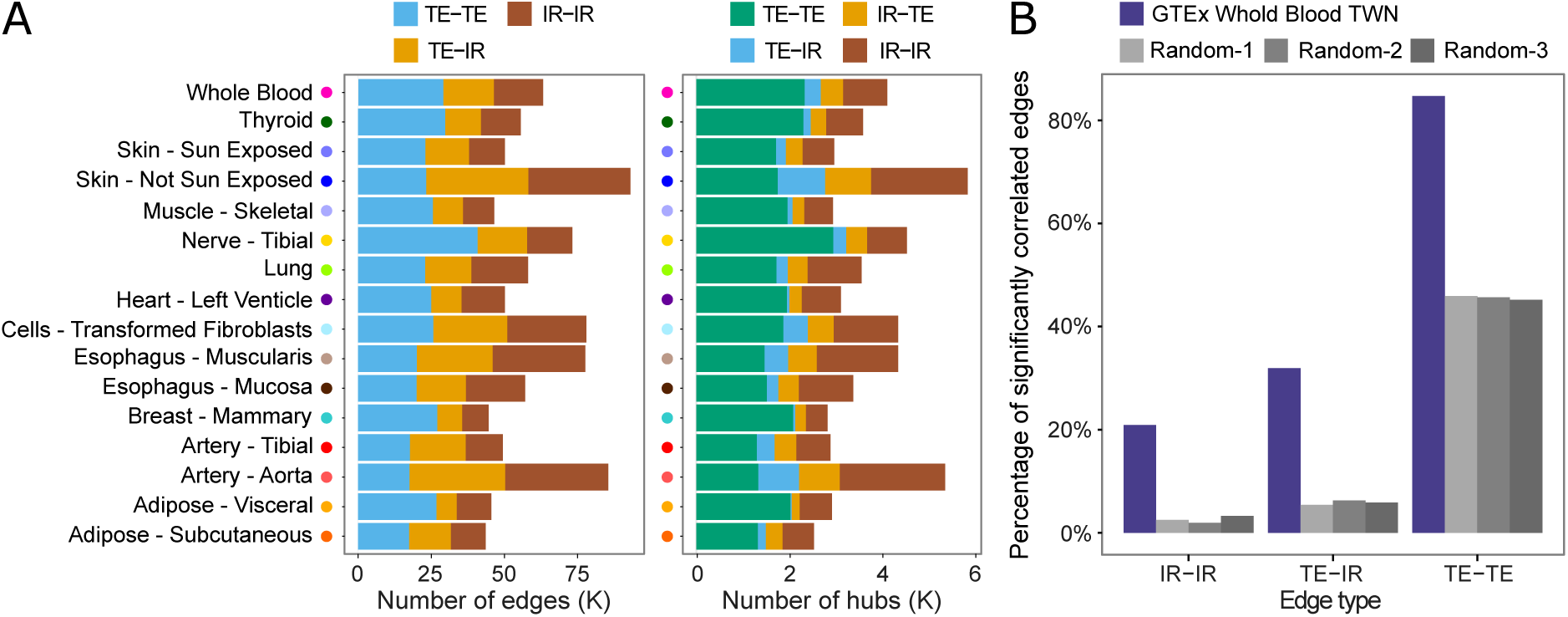
GTEx Transcriptome-Wide Networks summary and replication. A) For each tissue, number of edges and number of hub nodes (≥ 10 neighbors), segmented by the type of nodes connected by each edge (total expression or isoform ratio). For instance, a “TE-IR” hub is a total expression node with multiple isoform ratio neighbors, whereas a “IR-TE” hub is an isoform ratio node with multiple total expression neighbors. B) Fraction of whole blood TWN edges replicating in an independent RNA-seq sample (DGN) (Battle et al., 2014; Mostafavi et al., 2014).

Reconstructing co-expression networks requires estimation of a large number of parameters (in our case, over 2 × 10^8^) despite a small number of samples (≤ 430); robustness and replicability of network edges are thus important considerations. While there are not other large-scale RNA-sequencing data sets for most GTEx tissues, we replicated relationships identified by our GTEx whole blood TWN using an independent whole blood RNA-seq data set on 922 individuals of European ancestry, the Depression Genes and Networks study (DGN) (Battle et al., 2014; Mostafavi et al., 2014). First, we tested whether expression levels and/or isoform ratios directly connected by an edge in the GTEx whole blood TWN were also correlated in DGN. For all edge types, we found that a much higher fraction of the pairs connected by an edge in the GTEx TWN were significantly correlated in DGN compared to those from random networks (84.7% versus 45.6%, 31.9% versus 5.9%, and 20.9% versus 2.6% for TE-TE, TE-IR and IR-IR edges, respectively; FDR ≤ 0.05; Fig. 2B). Next, we constructed a TWN from DGN data over genes and isoforms common in both data sets. All pairs of nodes connected directly or indirectly in the GTEx whole blood TWN had significantly shorter network path distances in the DGN network compared to the distance in the same networks with the node labels shuffled (Wilcoxon rank-sum test, *p* ≤ 2.2 × 10^−16^; Supplementary Fig. 3). Together, these results show that, for each category of relationship identified in the GTEx TWN, including connections among isoform ratio nodes, we found support for shared relationships in DGN. This provides replication in an independent sample for the same tissue, despite different alignment and isoform quantification pipelines between the two data sets.

**Figure 3.**
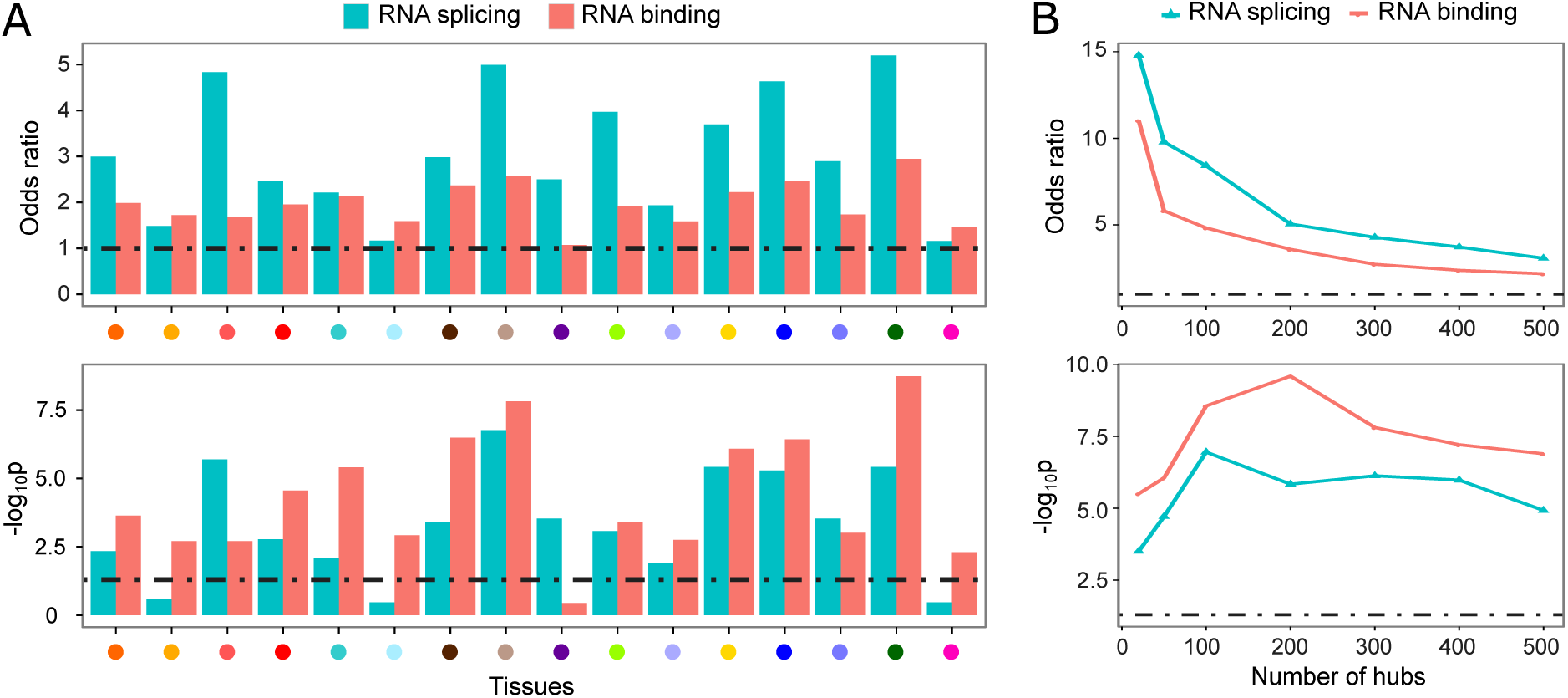
Enrichment of candidate splicing regulators among TWN hubs. A) In each per-tissue TWN, the odds ratio and p-value of enrichment among the top 500 TE-IR hub genes for GO annotations reflecting RNA binding and RNA splicing. B) Among consensus TE-IR hubs across all tissues, enrichment for GO annotations reflects RNA binding and RNA splicing functions.

### TWN hubs are enriched for regulators of splicing

We used the sixteen TWNs to characterize the regulation of relative isoform abundance in each GTEx tissue. Here, we focused on evaluation of network hubs. *Hub genes*, or genes with a high number of neighbors, in a biological network tend to be essential in biological mechanisms and, in co-expression networks, are more likely to have regulatory function (Barabasi and Oltvai, 2004; Jeong et al., 2001; Albert, 2005). Unlike traditional networks, TWNs have two different types of nodes—total expression and isoform ratio—therefore, we have four categories of hub genes that likely reflect different essential and regulatory function (Fig. 1C). For instance, a transcription factor may appear as a total expression node with many total expression neighbors. On the other hand, a hub arising from a total expression node connected to a large number of isoform ratio nodes may reflect a gene important in regulation of alternative splicing. We identified the top hub nodes by degree centrality for all categories in each of the 16 tissues (Supplementary Table 2). To avoid bias due to different numbers of isoforms per gene, we measured degree centrality of a node by the number of unique genes among neighboring nodes in each category (Methods).

We investigated whether hub nodes with many isoform ratio neighbors were likely to be regulators of alternative splicing. For each tissue, we evaluated the top TE-IR hubs for enrichment of Gene Ontology (GO) terms related to RNA splicing, and observed a significant abundance of known RNA splicing genes (annotated with GO:0008380) among the top TE-IR hubs. Indeed, 13 of 16 tissues (81.25%) showed significant enrichment of RNA splicing genes in the top 500 TE-IR hubs (significance assessed at BH corrected *p* ≤ 0.05; median across all tissues *p* ≤ 6.22 × 10^−4^, Fisher’s exact test), and every tissue had larger than unit odds ratio of RNA splicing genes among the top hubs (Fig. 3A). Enrichment was robust to choice of hub degree threshold (Supplementary Fig. 4). Next, we tested for enrichment of RNA-binding proteins, many of which are known to be important regulators of RNA splicing and processing (Witten and Ule, 2011; Wang and Burge, 2008; Chen and Manley, 2009). We found that RNA binding genes (annotated with GO:0003723) were also significantly enriched, at Benjamini-Hochberg (BH) corrected *p* ≤ 0.05, among the top TE-IR hubs of every tissue except *heart – left ventricle* (median *p* ≤ 3.17×10^−4^; Fig 3A). Consider all GO terms, splicing, RNA binding, and RNA processing terms were consistently among the most enriched for TE-IR hubs across tissues (Supplementary Tables 3,4). The replication network estimated from the DGN data also indicated enrichment of both RNA splicing and RNA binding genes among TE-IR hubs (RNA splicing: *p* ≤ 1.7 ×10^−5^, odds ratio 2.72; RNA binding: *p* ≤ 2.5 ×10^−11^, odds ratio 2.37).

**Figure 4.**
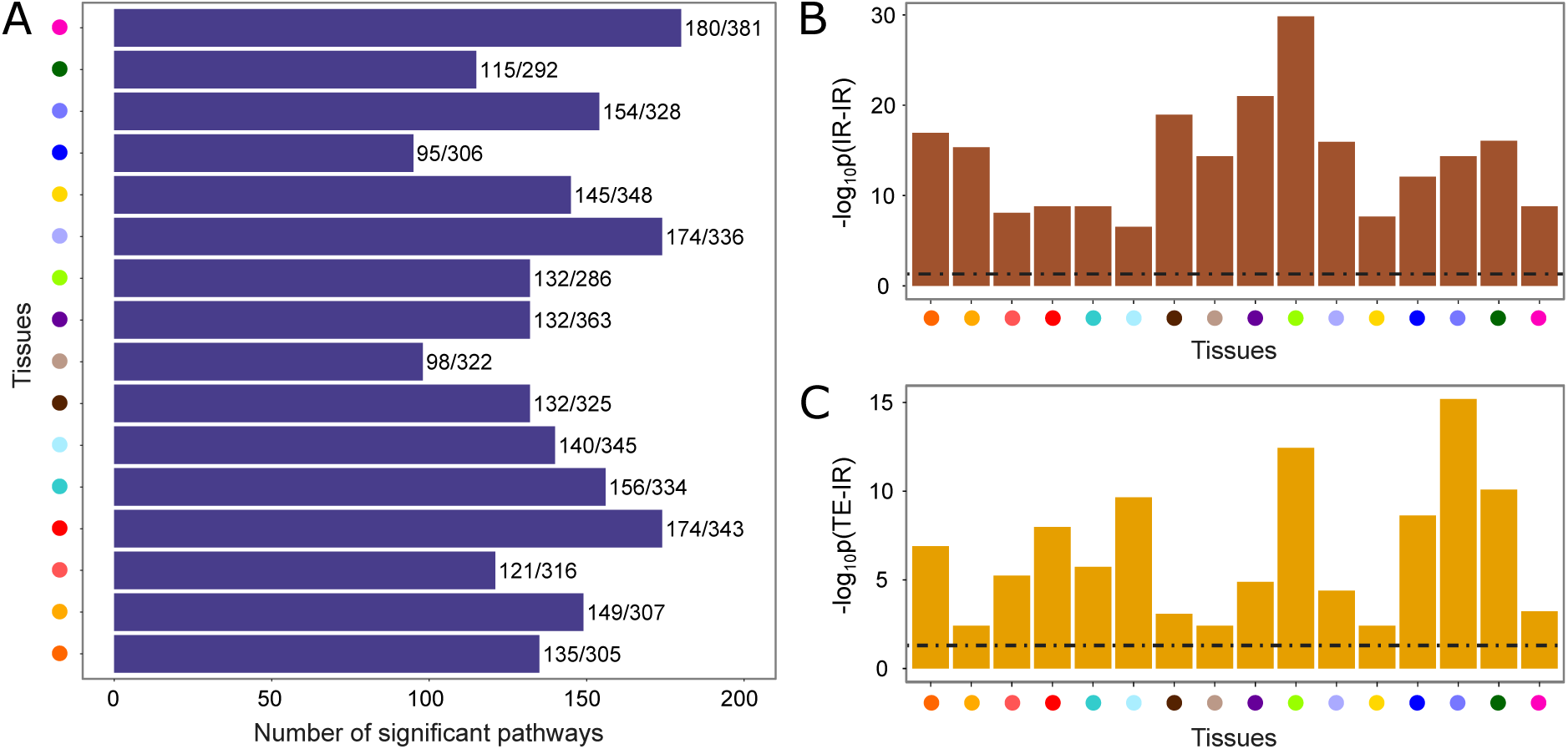
Pathway enrichment in TWNs. The colors representing the tissues are matched with tissue names in Fig. 1. (A) Per-tissue, the number of Reactome pathways enriched among connected components / total number of tested pathways for that tissue, considering only total expression nodes. (B) Enrichment for shared Reactome pathway annotation among gene pairs connected by edges between two isoform ratio nodes. (C) Enrichment for shared Reactome pathway annotation among gene pairs connected by an edge between a total expression node and an isoform ratio node.

Many regulatory relationships are in fact shared between tissues, and assessing hubs across all tissues jointly may improve robustness to noise in network reconstruction from limited data (Ballouz et al., 2015). Therefore, we identified TE-IR hubs shared across tissues (Table 1) using rank-product (Zhong et al., 2014). We first assigned lower ranks to hub genes with higher numbers of neighbors in each network, then aggregated the ranks of those genes across all networks by computing the product of these ranks. We sorted genes by rank product to find the top TE-IR hubs, having the largest number of neighbors in the most tissues (Methods). We again tested for enrichment of RNA splicing and RNA binding genes. Both p-values (Fisher’s exact test) and odds ratios were noticeably stronger in the joint analysis than in individual tissues; for example, we observed odds ratios of 9.8 for RNA splicing and 5.8 for RNA binding among the top 50 shared hubs (Fig. 3B).

**Table 1.** Top 20 cross-tissue TE-IR hubs. *Rank* is the rank-product rank of the gene. #Tissues is the number of tissues, out of 16, for which the hub gene (total expression) has at least one isoform ratio neighbor.

| Rank | Hub gene | #Tissues | Evidence |
| --- | --- | --- | --- |
| 1 | *TMEM160* | 16 |  |
| 2 | *RBM14* | 15 | Nuclear receptor coactivator that interacts with *TRBP* to regulate splicing in a promoter-dependent manner. (Auboeuf et al., 2004, 2002; Sui et al., 2007) |
| 3 | *ZMAT1* | 16 |  |
| 4 | *PPP1R10* | 15 | Mass spectrometry analysis suggests its involvement in pre-mRNA splicing through interaction with *ZNF638* (Du et al., 2014) |
| 5 | *ODC1* | 16 |  |
| 6 | *MGEA5* | 16 |  |
| 7 | *KLHL9* | 14 |  |
| 8 | *SRRM2* | 15 | Helps forming large splicing enhancing complexes (Chen and Manley, 2009). A mutation in *SRRM2* predisposes papillary thyroid carcinoma by changing alternative splicing (Tomsic et al., 2015). |
| 9 | *SRSF11* | 14 | A known serine/arginine-rich splicing factor (Zhang and Wu, 1996; Wu et al., 2006) |
| 10 | *ZNF692* | 15 |  |
| 11 | *ARGLU1* | 16 | Arginine/glutamate rich gene modulates splicing affecting neurodevelopmental defects (Magomedova et al., 2016). |
| 12 | *PPRC1* | 16 | Encodes protein similar to *PPARGC1* that regulates multiple splicing events (Martínez-Redondo et al., 2016). |
| 13 | *LUC7L3* | 15 | Regulates splice-site selection (Zhou et al., 2008) and affects cardiac sodium channel splicing regulation. (Gao et al., 2013) |
| 14 | *DUSP1* | 16 |  |
| 15 | *FOSL2* | 16 |  |
| 16 | *XPO1* | 16 | Interacts with *TBX3* (Kulisz and Simon, 2008) that regulates alternative splicing in vivo (mouse). (Kumar P. et al., 2014) |
| 17 | *PNISR* | 15 | Interacts with *PNN*, a suggested splicing regulator, and colocalizes with *Srrp300*, a known component of the splicing machinery (Zimowska et al., 2003). |
| 18 | *PNN* | 12 | Likely to be involved in RNA metabolism including splicing (Li et al., 2003) |
| 19 | *PTMS* | 12 | Involved in RNA synthesis processing (Vareli et al., 2000). |
| 20 | *CCDC85B* | 15 |  |

Many of the top ranked TE-IR hubs shared across tissues are indeed known to regulate splicing. *RBM14* (rank two), a RNA binding gene also known as *CoAA*, interacts with a transcription regulator *TRBP* to regulate splicing in a promoter-dependent manner (Auboeuf et al., 2004, 2002). Another RNA binding gene *PPP1R10* (rank four) has been implicated in pre-mRNA splicing using mass spectrometry analysis (Du et al., 2014). *SRRM2* (rank eight) and *SRSF11* (rank nine) are also known splicing regulators (Chen and Manley, 2009; Blencowe et al., 2000; Zhang and Wu, 1996; Wu et al., 2006). For eleven of the top twenty cross-tissue TE-IR hubs, we found previous work supporting a role in the regulation of splicing (Table 1). Together, these results suggested that TWN hubs are informative of splicing regulation and consistent with previous results showing that RNA binding proteins are principal splicing regulators. Uncharacterized TE-IR hub genes in a TWN are good candidates for investigating potential effects on splicing and relative isoform abundance.

### Co-regulation of expression and isoform ratios reflect biological pathways

It has been observed that genes with similar function or that participate in the same pathway often have correlated patterns of gene expression (Khatri et al., 2012; Hormozdiari et al., 2015). In the GTEx TWNs, we observed enrichment of edges between transcription factors and known target genes (Methods, Supplementary Fig. 5), as expected (Prieto et al., 2008; Roider et al., 2009). We also observed enrichment of closely connected genes for a number of Reactome pathways (Fabregat et al., 2016) as compared with permuted networks (95 − 180 pathways enriched per tissue at Bonferroni corrected *p* ≤ 0.05, Wilcoxon rank-sum test; Fig. 4A, Methods).

**Figure 5.**
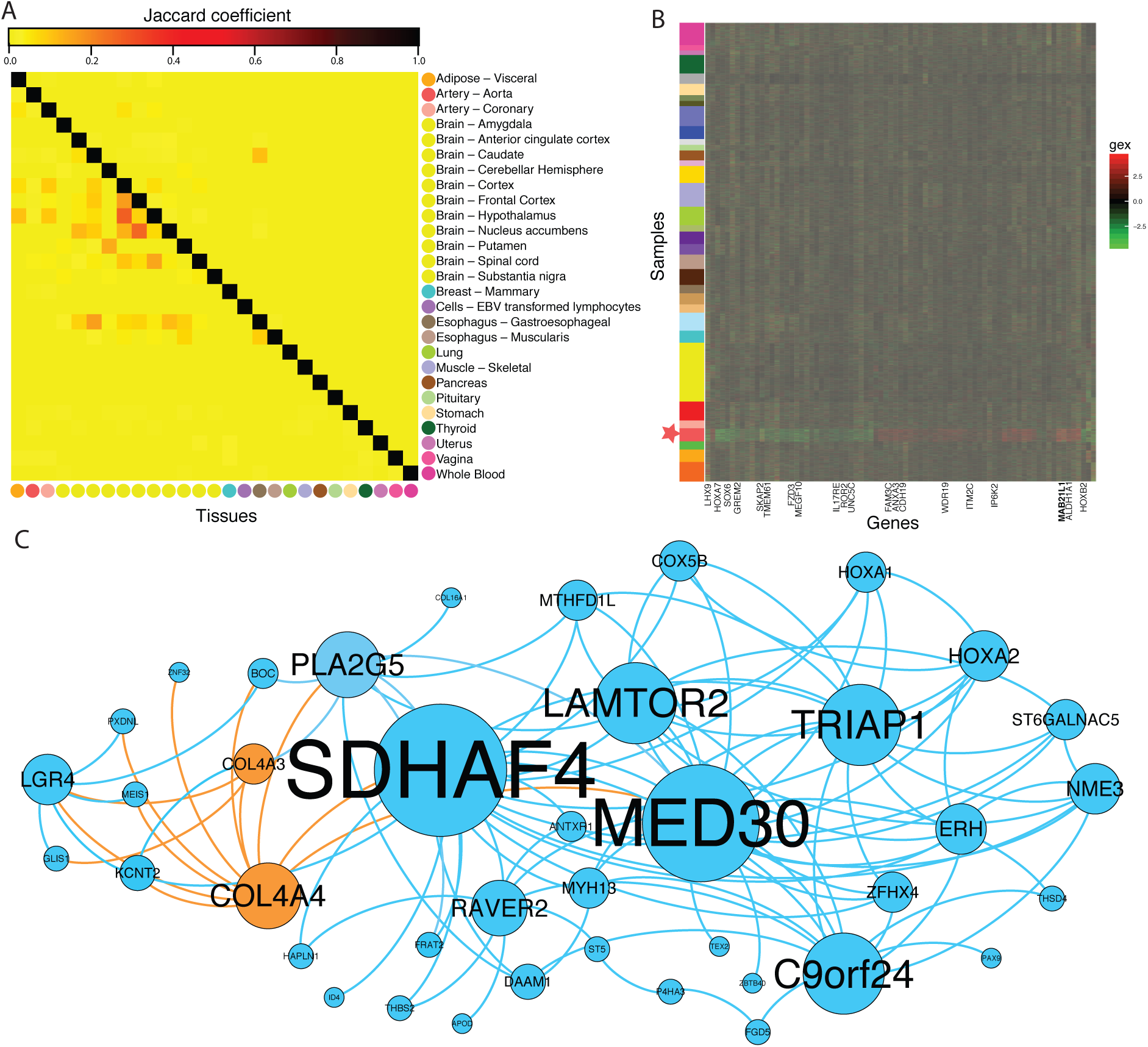
Cross tissue comparison of TSN results. (A) Jaccard coefficient quantified on shared edges (upper triangular) and shared nodes (lower triangular) across pairs of TSNs; (B) Gene expression levels, removing factors from BicMix not included in the network, for the genes identified in the TSN for *artery – aorta*. The y-axis is ordered by similarity to *artery – aorta*, with a star by the samples from *artery – aorta*. The colors on the y-axis correspond to the GTEx tissue legend above. The x-axis is ordered by expression similarity (i.e., hierarchical clustering), and hub genes are labeled with the large hub denoted in bold; (C) tissue-specific network for *artery – aorta*. Node size reflects betweenness centrality of the nodes. Orange nodes reflect replication in the BioCarta *acute myocardial infarction* (AMI) pathway; orange edges show the neighbors of the AMI pathway nodes.

In contrast to correlations among total expression levels of genes, patterns of correlation among relative isoform abundances are far less well-studied, and it has not been established whether or not the regulation of splicing is coordinated across functionally related genes. Initial studies have identified such correlation in particular tissues (Iancu et al., 2015) and specific processes (Dai et al., 2012). We evaluated each TWN for enrichment of edges between functionally related genes. For all 16 tissues, the TWNs demonstrated significant abundance of edges between isoform ratios of two distinct genes that participate in the same Reactome pathway (all tissues significant at BH corrected *p* ≤ 0.05; median *p* ≤ 10^−14^; Fig. 4A). Similarly, edges between total expression of one gene and isoform ratio of a second gene were enriched for a gene annotation in the same pathway (median *p* ≤ 10^−5^; Fig. 4B). As expected, we also observed pathway enrichment for edges between total expression nodes (Supplementary Fig. 6). The patterns of functional enrichment were in fact somewhat stronger among total expression nodes, which could be influenced by more accurate quantification of total expression compared with isoform ratios from RNA-seq data, derivation of functional annotation from gene expression studies, or tighter co-regulation of transcription compared with splicing of functionally related genes. These results provide evidence in a large, multitissue data set that functionally related genes display correlated variation in relative isoform abundance, indicative of co-regulation of splicing. Leveraging this finding allows TWNs to be used to identify sets of possibly co-functional genes, or to predict the function of unknown genes (Warde-Farley et al., 2010) based on a more comprehensive understanding of co-regulation including regulation of splicing.

### Comparison between TWNs reveals per-tissue hub genes

We evaluated the overall similarity of the TWNs between tissues. We tested concordance of hubs between each pair of tissues using Kendall’s rank correlation computed over genes ordered by degree centrality (Supplementary Fig. 7). We observed greater than random levels of similarity between most tissues for all hub types (median *p* ≤ 1.0 × 10^−5^ for each hub type) and functionally related tissues displaying greater levels of similarity. For example, the two skin tissues were grouped together for each hub type, and were found to be similar to *esophagus – mucosa*, which contains primarily epithelial tissue (Squier and Kremer, 2001). *Skeletal muscle* and *heart – left ventricle* grouped together, and *breast* was highly similar to the two adipose tissues, reflecting shared adipose cell type composition. While these results may be partially influenced by overlap in donors, they provide evidence that splicing displays variability between tissues and is regulated in a tissue-specific manner (Qian et al., 2005; Ong and Corces, 2011).

To identify candidate tissue-specific regulatory genes, we evaluated TE-IR hubs that were ranked high by degree centrality in related tissues, but ranked low among unrelated tissues (Methods, Supplementary Table 5). Several of the top ranked tissue-specific hubs were genes with evidence of known tissue-specific function or relevance. In the tissue group including *breast* and two adipose tissues, the top tissue-specific TE-IR hub was *TTC36*, a gene highly expressed in breast cancer (Liu et al., 2008). The top ranked gene hub for the *skeletal muscle* and *heart – left ventricle* tissues group was *PLEKHG4*, which has been reported to be associated with cardiac failure in mice (Mullick et al., 2006).

### Tissue-specific networks identify gene co-expression patterns unique to tissues

To directly assess the tissue-specificity of co-expression relationships, we built tissue=specific networks by considering all GTEx samples across tissues simultaneously, decomposing co-expression signal into co-expression signals shared across tissues and those specific to individual tissues. To do this, we applied a Bayesian biclustering framework, BicMix, and reconstructed tissue-specific networks from the fitted model (Gao et al., 2016). BicMix uses a sparsity-inducing prior to differentiate between co-expression relationships specific to a single tissue and those shared across tissues, controlling for batch effects, population structure, and shared individual effects across tissues (Gao et al., 2016). The strength of this joint approach applied to over 7,000 RNA-seq samples is that, with more thorough and comprehensive sampling of heterogeneous tissues types, we are able to isolate co-expression signals unique to single tissues and to reconstruct much more precise tissue-specific networks (TSNs).

We built TSNs for 27 GTEx tissues. Here, we limited network nodes to total gene expression rather than isoform ratio for simplicity. Across the 27 TSNs, the mean number of nodes (considering only genes with tissue-specific edges) was 48, and the average number of edges was 436. However, this average is driven by four larger networks (Supplementary Table 6; Supplementary Fig. 8) with over 100 nodes each: *thyroid* (216), *stomach* (179), *whole blood* (174), and *vagina* (101). There are many fewer nodes in these TSNs as compared to the TWNs because we only included genes in the network that had one or more tissue-specific edge. For the eight tissues for which we also constructed TWNs, we found that tissue-specific networks indeed overlapped with the TWNs, generally showing greater concordance for the matched tissue (Supplementary Fig. 9). As expected, TSNs contained a small subset of edges from full per-tissue TWN, representing the co-expression components that are tissue-specific rather than shared.

Additionally, we built four tissue-specific networks for subsets of similar tissues (GTEx Consortium, 2015) to capture gene relationships common within each subset but unique compared with all other tissues. Most tissues showed expression patterns close to at least one other assayed tissue (Supplementary Fig. 10), leading to a depletion of tissue-specific effects, and motivating evaluation of similar tissues together. We considered four groups: 1) all 13 brain tissues; 2) two adipose tissues and *breast*; 3) two heart tissues and three artery tissues and 4) four digestive tissues. On average, these tissue subset networks contained 4,779 edges and 189 nodes. However, this was skewed by the brain network, which contained 18,854 edges connecting 648 nodes. Excluding the brain network, we found 87 edges and 36 nodes on average across the other three tissue subset networks. The 27 single tissue TSNs and the four tissue subset networks are available at http://gtexportal.org (in progress).

#### Functional analysis of tissue-specific networks

We investigated the functional properties of the genes and edges identified in each tissue-specific network. First, we measured sharing of network components between the 27 distinct TSNs. We found minimal sharing of network nodes and even less sharing of network edges among all pairs of tissues (Jaccard coefficient; Fig. 5A). This was expected as a result of BicMix’s strong control over shared individuals and gene expression levels across tissues. Tissue pairs that appeared to share genes predominantly included brain tissues. We confirmed enrichment of known tissue-specific genes in the TSNs using a previously defined list of Gene Ontology (GO) terms (Ashburner et al., 2000) indicative of tissue-specific function available for eleven tissues (Pierson et al., 2015). We found four TSNs nominally enriched for genes with specificity in the matched tissue, namely *artery – coronary* (B-H corrected *p* ≤ 0.23), *EBV transformed lymphoblastoid cell lines* (with blood, B-H corrected *p* ≤ 0.09), *skeletal muscle* (B-H corrected *p* ≤ 0.13), and *stomach* (B-H corrected *p* ≤ 0.13). Significant cross tissue enrichments were observed in a small number of tissues. In the *vagina* TSN, pituitary genes were significantly enriched (B-H corrected *p* ≤ 4.35 × 10^−5^) and in the *artery – aorta* TSN, pituitary genes were significantly enriched (B-H corrected *p* ≤ 0.0049).

Next, we evaluated the hub genes in each TSN, considering three thresholds of centrality: ≥ 5 edges (“small hubs”), ≥ 10 edges (“hubs”), and ≥ 50 edges (“large hubs”). Hubs were not enriched overall for cross-tissue transcription factors (TFs) (hypergeometric test across all TSNs, *p* ≤ 0.84; small and large hubs showed similar results), or for cross-tissue and tissue-specific TFs (hypergeometric test across all TSNs, *p* ≤ 0.90; small and large hubs showed similar results); this result echos previous work on transcription factor enrichment in genes with cis-eQTLs that appear to have broad regulatory effects on transcription (Jo et al., 2016; Weiser et al., 2014). However, hubs in several networks included genes known to play a role in tissue-specific function and disease. Specifically, we found that the single large hub in *brain – caudate*, *MAGOH*, which is a part of the exon junction complex that binds RNA, has been found to regulate brain size in mice through its role in neural stem cell division (Silver et al., 2010). The single large hub for *artery – aorta*, *MAB21L1*, has been shown to be an essential gene for embryonic heart and liver development in mice by regulating cell proliferation of proepicardial cells (Saito et al., 2012).

Additionally, we measured enrichment of known pathways in the TSNs. While we did not observe enrichment across all tissues, we found that the *hematopoietic cell lineage* KEGG pathway was significantly enriched in the TSN for *EBV transformed lymphoblastoid cell lines* (*LCLs*; Fisher’s test, B-H adjusted *p* ≤ 0.05); a hematopoietic stem cell is the developmental precursor of leukocytes, reflecting the expression signature of the parent cell type. The *LCL* TSN also had significant enrichment in the BioCarta *IL-17 signaling* and *T cytotoxic cell surface molecules* pathways (LCLs; Fisher’s test, B-H adjusted *p* ≤ 1.50 × 10^−4^). *IL-17* is a cytokine produced from T-cells that is involved in inflammation, and the cytotoxic t-cell pathway is involved in eliminating cells with certain surface antibodies. Although not significant after multiple testing correction, *artery – coronary* showed enrichment in four tissue-relevant pathways (uncorrected *p* ≤ 0.016 for all): the *ACE2* pathway, which regulates heart function, *acute myocardial infarction* (AMI) pathway, the *intrinsic prothrombin activation* pathway, which is involved in one phase of blood coagulation, the *platelet amyloid precursor protein (APP)* pathway, which includes genes involved in anti-coagulation functions in platelets and senile plaques, and the *vitamin C in the brain* pathway, which is responsible for the cellular uptake of reduced ascorbate from platelets (Fig. 5).

### Integration of networks with regulatory genetic variants

Both TWNs and TSNs were estimated using gene expression data alone. However, the GTEx v6 data also include genotype information for each donor. We intersected the edges detected by our networks with QTL (quantitative trait locus) association statistics to replicate specific network edges through evidence of conditional associations with genetic variants across those edges and to increase power to detect long range (trans) effects of genetic variation on gene regulation. First, we demonstrated that, for both TWNs and TSNs, there was enrichment for associations between the top cis-eVariant (the variant with lowest p-value per gene with a significant cis-eQTL) for each gene and the expression level or isoform ratio of its network neighbors based on QTL mapping in the corresponding tissue (Fig. 6). This provides evidence of a relationship between the pairs of genes connected by an edge. For TWNs, among total expression nodes with an isoform ratio neighbor, we found evidence for 61 trans (i.e., inter-chromosomal) associations and 86 intra-chromosomal associations tested between a cis-eVariant for the total expression gene and the isoform ratio of the neighboring node (FDR ≤ 0.05), providing support for these TWN edges. Our top two associations were between two variants, rs113305055 in *artery – tibial* and rs59153288 in *breast* (both near *TMEM160*), with isoform ratios of *CST3* (*p* ≤ 9.3 × 10^−8^, and *p* ≤ 4.0 × 10^−7^, respectively). *TMEM160* is the top cross-tissue hub in our TWNs with many IR neighbors (Table 1). Thus, we tested for association of these variants with all isoform ratios genome-wide and observed a substantial enrichment of low p-values in numerous tissues (Fig 6A; Supplementary Fig. 11). In the TSNs, we identified seven cis-eVariants across five tissues associated with eight different trans-eGenes through seven unique cis-eGene targets, one of which was intra-chromosomal (FDR ≤ 0.2; Supplementary Table 7). We also observed enrichment for low p-values over the tests corresponding to each network edge (Fig. 6B).

**Figure 6.**
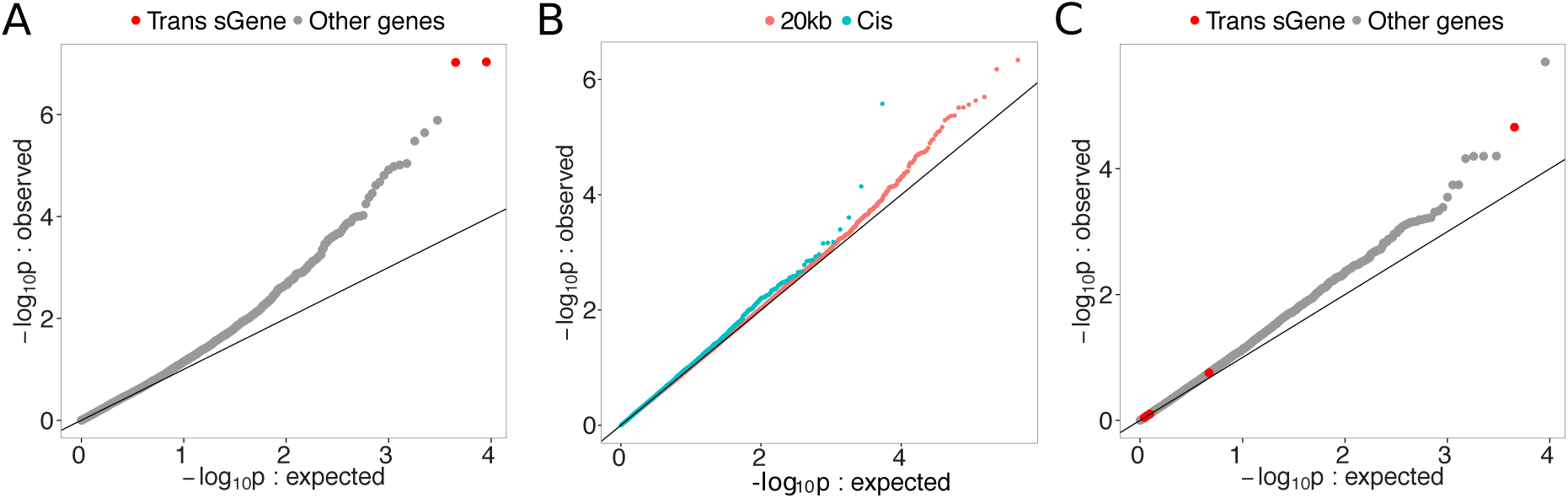
Association of local genetic variants with distant network neighbors. (A) Enrichment of association between rs113305055, a genetic variant near a cross-tissue TWN hub *TMEM160*, with all isoform ratios genome-wide in *artery – tibial*. (B) Enrichment of associations between local genetic variants (either the top cis-eVariant or any variant with 20 Kb) of each gene, and network neighbors in the tissue-specific networks. (C) Enrichment of association between rs115419420, a genetic variant local to *CRELD1*, with all isoform ratios in *skeletal muscle*.

We used the TWNs and TSNs in a trans-QTL discovery framework that does not rely on prior cis-eQTL association testing. In the test described above, it is possible that artifactual correlations among gene expression levels, identified as edges in the networks, may also lead to artifactual trans-eQTL associations among cis-eVariants and the neighboring genes of the cis-eGene detected from the same data. However, large-scale, independent RNA-seq data sets are unavailable for most of the tissue types represented in GTEx, and identifying trans-eQTLs from a standard genome-wide association test has proven challenging (Jo et al., 2016), in part due to the large number of statistical tests. Restricting association tests to a plausible subset, according to prior knowledge (Jo et al., 2016) such as network relationships (Weiser et al., 2014) can increase statistical power substantially.

Here, we identified novel trans-splicing QTLs (trans-sQTLs) and trans-expression QTLs (trans-eQTLs) by restricting the set of tests to those suggested by the TWNs and TSN edges in the corresponding tissue. From the TWNs, we sought to identify trans-splicing QTLs (sQTLs) based on TE-IR hub genes, using the top 500 hubs by degree centrality. We tested every SNP within 20 Kb of the TE hub-gene’s transcription start site (TSS) for association with isoform ratios of each neighbor in the TWN. Using this approach, we identified 58 trans-sQTLs corresponding to six unique genes (sGenes) at FDR ≤ 0.2 (Table 2). For example, we identified a trans-sQTL association in *skeletal muscle* between rs115419420 and *CARNS1* (*p* ≤ 2.18 × 10^−5^) that is supported by a cis association with the TE-IR hub *CRELD1*. This eVariant also showed enrichment for low p-values with numerous isoform ratios genome wide (Fig. 6C). In the TSNs, we identified 27 trans-eQTLs using variants within 20 Kb of each gene and testing for association with the neighbors of those genes in the gene expression data of the same tissue (FDR ≤ 0.2; Supplementary Table 8). All of these associations were with inter-chromosomal. Overall, we saw a substantial enrichment of p-values for association between genetic variants local to a gene and the genes’ neighbors in each network (Fig. 6B).

**Table 2.** Top 20 cross-tissue TE-IR hubs. P-value and FDR for association between the variant and the trans-sGene listed. Local gene target listed for reference.

| variant | trans-eTranscript | trans-sGene | local gene | p-value | FDR | tissue |
| --- | --- | --- | --- | --- | --- | --- |
| rs6122466 | ENST00000496440.1 | *CEP350* | *PPDP* | $9.08 \times 10^{-7}$ | 0.09 | Adipose – Visceral |
| rs397828484 | ENST00000528430.1 | *PPP1R16A* | *NMRK2* | $1.66 \times 10^{-6}$ | 0.10 | Muscle – Skeletal |
| rs7668429 | ENST00000340875.5 | *MEF2D* | *CLOCK* | $4.81 \times 10^{-6}$ | 0.10 | Muscle – Skeletal |
| rs7980880 | ENST00000409273.1 | *XIRP2* | *CALCOCO1* | $9.91 \times 10^{-6}$ | 0.11 | Muscle – Skeletal |
| rs56359342 | ENST00000396435.3 | *IQSEC2* | *CRAMP1L* | $1.43 \times 10^{-5}$ | 0.14 | Muscle – Skeletal |
| rs115419420 | ENST00000531388.1 | *CARNS1* | *CRELD1* | $2.18 \times 10^{-5}$ | 0.19 | Muscle – Skeletal |

## Discussion

We reconstructed co-expression networks that capture novel regulatory relationships in diverse human tissues using large-scale RNA-seq data from the GTEx project. First, we specified an approach for integrating both total expression and relative isoform ratios in a single sparse Transcriptome-Wide Network (TWN) based on graphical lasso. Splicing is a critical process in a number of tissue- and disease-specific processes and pathways (Ghigna et al., 2008; Hutton et al., 1998; Glatz et al., 2006; D’Souza et al., 1999), but, critically, isoform ratios have not been included in co-expression network analysis to allow the study of splicing regulation. We estimated TWNs from sixteen tissues and demonstrated that hubs in TWNs are strongly enriched for genes involved in RNA binding and RNA splicing. We found that, across tissues, the top hub genes with isoform ratio neighbors included many genes with known impact on splicing such as *RBM14*, a hub in all 16 tissues with TWNs. We identified a number of novel shared and tissue-specific candidate regulators of alternative splicing. TWNs may be extended to include diverse expression phenotypes such as allelic expression or individual splicing event frequencies. As more large-scale RNA-seq studies become available, this method will be useful for analyzing diverse types of regulatory relationships in disease, longitudinal, and context-specific studies.

Next, we estimated tissue-specific networks for 27 single tissues and across four tissue subsets; these networks represent co-expression relationships unique to individual tissues and sets of closely related tissues. Distinguishing between shared and tissue-specific structure across single tissue co-expression networks is challenging, but is essential for understanding tissue-specific regulatory processes in disease. From these tissue-specific networks, we identified hub genes involved in the essential tissue-specific regulation of transcription, such as *MAGOH* in the *brain – caudate* specific network and *MAB21L1* in the *artery – aorta* specific network, both of which are essential for the development of their specific organs. We used these networks to quantify shared relationships across tissues, and found minimal sharing of relationships across these 27 tissues. Finally, we replicated edges in our networks using integration of genetic variation, and we identified 33 novel trans-QTLs affecting both expression and splicing. Together, our results provide the most comprehensive map of gene regulation, splicing, and co-expression in the largest set of tissues available to date. These networks will provide a basis for interpreting the transcriptome-wide effects of genetic variation, differential expression and splicing in complex disease, and impact of diverse regulatory genes in the human genome.

## Methods

### RNA-seq data from GTEx project data

The NIH Common Fund’s Genotype-Tissue Expression (GTEx) consortium (GTEx Consortium, 2015) provides RNA-seq and microarray experiments. The original GTEx RNA-seq samples have been obtained from recently deceased donors (samples collected within 24 hours), between ages 21 and 70, BMI 18.5 to 35, and not under exclusionary medical criteria such as whole-blood transfusion within 24 hours or infection with HIV. The blood samples have been extracted for both genotyping with Illumina HumanOmni 2.5M and 5M BeadChips, as well as EBV-transformation of lymphoblastoids into cell lines. Then, biopsies from a set of tissues from different body sites (averaging about 28 per individual) have been obtained, stabilized with PAXgene Tissue kits, and then shipped to designated facilities for paraffin embedding, sectioning, and analyses of histology. After quality control protocols that checked for evidence of autolysis, inflammation and other pathology that could affect RNA-seq results, the stabilized tissue samples were sent to sequencing facilities for DNA/RNA extraction from the samples and performing both microarray and RNA-seq experiments. In particular, the RNA-seq experiments were performed with Illumina Hi-Seq 2000 following the TrueSeq RNA protocol, yielding 76-bp paired-end reads averaging approximately 50 million reads per sample. As a result, we have 8,551 total experiments from 449 individuals for phase 1.

### RNA-sequencing alignment and transcript quantification

The RNA-seq processing pipeline follows previously described steps (McDowell et al., 2016). Adapter sequences and overrepresented contaminant sequences, identified by FastQC (v.0.10.1) (Andrews, 2010), were trimmed using Trimmomatic (v.0.30) (Bolger et al., 2014) with 2 seed mismatches and a simple clip threshold of 20. Leading and trailing nucleotides (low quality or *N*s) were trimmed from all reads until a canonical base was encountered with quality greater than 3. For adaptive quality trimming, reads were scanned with a 4-base sliding window, trimming when the average quality per base dropped below 20. Any remaining sequences shorter than 30 nucleotides were discarded.

We aligned the RNA-seq reads using the Star aligner in 2-pass mode (Dobin et al., 2013). After preparing the genome with STAR aligner genomeGenerate mode using a splice junction database (sjdbGTFfile) set to GENCODE v.19 annotation, the splice junction database overhang (sjdbOverhang) set to 75 bp, and defaults for all remaining settings. STAR aligner alignReads mode was run using default settings except `outFilterMultimapNmax` was set to 1 so that only uniquely mapping reads were retained.

We performed transcript and gene quantification using RSEM v1.2.20 (Li and Dewey, 2011). We used default settings, allowing for four threads and using paired-end aware quantification.

### Pre-processing for per-tissue TWNs

We considered only protein-coding genes on the autosomes and X chromosome to construct TWNs in all tissues. We filtered genes and isoforms using a threshold of 10 samples with ≥ 1 TPM and ≥ 10 samples with ≥ 6 reads. We also filtered out genes where the Ensembl gene ID did not uniquely map to a single HGNC gene symbol. Isoform ratio was computed by using annotated isoforms in Gencode V19 annotation, and undefined ratios (0/0, when none of the isoforms was expressed) were imputed from the mean ratio per isoform across individuals. Each gene’s least abundant isoform was excluded to avoid linear dependency between isoform ratio values. We log-transformed the total expression data and standardized both total expression levels and isoform ratios. To correct hidden confounding factors, we applied HCP (Hidden covariates with prior) (Mostafavi et al., 2013), whose parameters were selected based on an external signal relevant to regulatory relationships. Namely, we selected parameters that produced maximal replication of an independent set of trans-eQTLs from meta-analysis of a large collection of independent whole blood studies (Westra et al., 2013). For both total expression levels and isoform ratios of genes in all tissues, the best HCP parameters (*k* = 10, *λ* =1, *σ*_1_ =5, *σ*_2_ = 1), which consistently reproduced a largest subset of the gold-standard trans-eQTLs in GTEx *whole blood* samples even when subsetting the number of samples, were used for correcting data. Finally, quantile-normalization to a standard normal distribution was applied.

To avoid spurious associations due to miss-mapped reads, we filtered out genes and isoforms with mappability score constraints (≥ 0.97). We first downloaded mappability scores of all 75-mers and 36-mers in the human reference genome (hg19) from the UCSC genome browser for exonic regions and untranslated regions (UTRs), respectively (accession: wgEncodeEH000318, wgEncodeEH00032) (Derrien et al., 2012). For each gene, we then measured mappability scores for either exonic regions or UTRs with corresponding *k*-mers matched to the regions and aggregated the mappability scores for two regions by computing their weighted average. The weights were proportional to the total length of exonic regions and UTRs.

We filtered out isoforms of a gene if the mean IR of a most dominant isoform was ≥ 0.95. In each tissue, we further reduced the number of features to 6, 000 genes and 9, 000 isoforms for computational tractability. To do so, we first considered genes or isoforms if > 10 samples have TPM > 2 or reads > 6. To obtain the final set of genes, we first considered the top 9, 000 genes based on their average expression levels and then selected the top 6, 000 highly variable genes across individuals. Similarly, to obtain the final set of isoforms, we first considered the 13, 500 genes with the highest expressed isoform levels on average. We reduced this to 11, 250 genes based on the entropy of isoform ratios across individuals, normalized by the maximum entropy possible with the same number of isoforms, and finally took the top 9, 000 most highly variable isoforms in terms of TPM values.

### Per tissue Transcriptome-Wide Networks (TWNs)

We built per-tissue Transcriptome-Wide Networks (TWNs) using scalable graphical lasso (Hsieh et al., 2011). We estimated a sparse precision matrix (Θ) by optimizing the following objective with Λ specifying different penalties for different types of edges: 
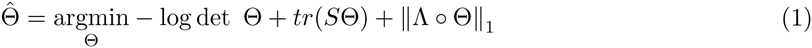
where the entry in *r*th row and *c*th column of Λ was

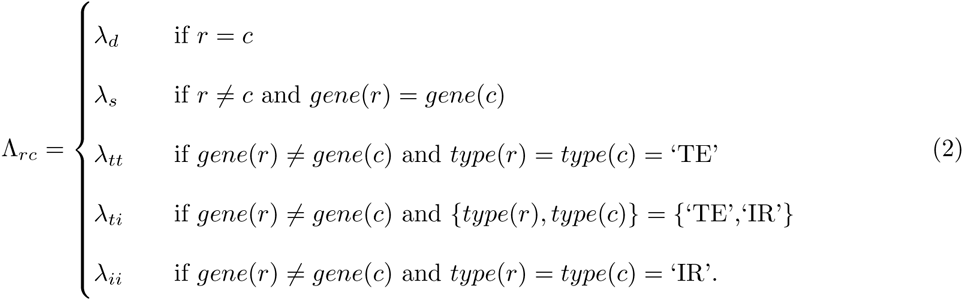

Here, *gene*(*k*) denotes the gene that the *k*th feature belongs to. *type*(*k*) denotes whether or not the *k*th feature represents total expression (‘TE’) or isoform ratio (‘IR’).

We did not penalize diagonal entries (*λ*_*d*_ = 0), and put a small non-zero penalty for edges between distinct features belonging to the same gene (*λ*_*s*_ =0.05), such as distinct isoforms of the same gene. We selected the other penalties (*λ*_*tt*_, *λ*_*ti*_, *λ*_*ii*_) such that the network has a scale-free topology with a reasonable number of edges. The empirical pairwise correlations distribution for different types of edges were different: correlations between two total expression nodes were generally much higher than those between two isoform ratio nodes or between a total expression node and an isoform ratio node (Supplementary Fig 1), while the later two distributions were apparently similar. We tried all (*λ*_*tt*_, *λ*_*ti*_, *λ*_*ii*_) combinations where *λ*_*tt*_*ɛ*{0.3, 0.35, 0.4, 0.45, 0.5}, *Λ*_*ti*_*ɛ*{0.25, 0.3, 0.35, 0.4}, and *λ*_*ti*_ = *λ*_*ii*_. We measured the scale-free property by the square of correlation (*R*^2^) between log(*p*(*d*)) and log(*d*), where *d* is an integer and *p*(*d*) represents the fraction of nodes in the network with *d* neighbors (Zhang and Horvath, 2005). We selected penalty parameters so that *R*^2^ ≈ 0.85 and there were at least 5,000 edges of each type. Selected parameters for each tissue are shown in Supplementary Table 1. Each non-zero element in Λ_*rc*_ in the precision matrix with selected penalty parameters represents an edge between the *r*th and *c*th features in our network.

We excluded some edges from our networks for quality purpose and interpretability. In other words, we excluded edges between nodes belonging to same gene for downstream analysis. Then, we aligned every 75-mer in exonic regions and 36-mers in UTRs of every gene with mappability *<* 1.0 to the reference human genome (hg19) using Bowtie (v 1.1.2) (Langmead et al., 2009). If any of the alignments started within an exon or a UTR of another gene, then these two genes were considered “cross-mappable”, and we excluded edges between these pairs of genes. We also excluded edges between genes with overlapping positions in the reference genome to avoid mapping artifacts.

### Replication of whole blood TWN

We replicated our network edges with GTEx whole blood tissue in an independent RNA-seq data set: Depression Genes and Networks (DGN) (Battle et al., 2014; Mostafavi et al., 2014). These RNA-seq data include 15,231 genes and 12,080 isoforms from whole blood in 922 samples, out of which 5,609 genes and 1,464 isoforms were uniquely mapped to the set of genes and isoforms used in GTEx *whole blood* network construction. Firstly, to check if the genes and isoforms directly connected in the GTEx *whole blood* network were supported by correlation in the DGN data set, we computed the fraction of significantly correlated (Spearman correlation, FDR ≤ 0.05) gene-gene/gene-isoform/isoform-isoform pairs in DGN. We then compared these fractions with those in random pairs generated by permuting genes/isoforms labels in the TWN. Next, to verify if our method could reproduce relationships in GTEx *whole blood* network from DGN data, we tested if node pairs connected directly or indirectly in the GTEx *whole blood* network had a shorter distance between them in the DGN network compared to the same network with the node labels shuffled. We performed a one-sided Wilcoxon rank-sum test between two groups: i) pairwise distances between GTEx-connected TE-TE / TE-IR / IR-IR pairs in the DGN network. ii) those in random DGN networks generated by permuting genes/isoforms among themselves. Here we generated random networks ten times to estimate the null distribution.

### Hub ranking

We ordered the network hubs by degree centrality for each tissue according to the number of unique gene-level connections to avoid the effect of different number of isoforms per gene. To do this, we created a gene-level network from original TWNs by keeping TE nodes as they were and grouping all isoforms of the same gene together to form a compound IR node. We put an edge between a compound IR node and a total expression node (or another compound isoform ratio node) if any isoform of the compound had an edge with the TE node (or any isoform of the other compound) in the original TWN, and the weight is equal to the sum of absolute weights of all such edges in the original TWN. TE-TE and IR-TE hubs were ordered by the number of TE nodes they were connected with. TE-IR and IR-IR hubs were ordered by the number of compound IR nodes they were connected with. If multiple hubs had the same number of connections, ties were broken by the sum of corresponding edge weights.

### Splicing and RNA binding enrichment in top TE-IR hubs

We downloaded a list of human genes annotated with *RNA Splicing* (GO:0008380) and *RNA Binding* (GO:0003723) using topGO (Alexa and Rahnenfuhrer, 2016). We computed the enrichment of these RNA splicing and RNA binding genes in top 500 TE-IR hubs using Fisher’s exact test. The set of all genes represented in the corresponding network was used as background.

### General hubs and tissue-specific hubs

We used rank-product (Zhong et al., 2014) to find hubs generally ranked high in a set of tissues. We first ranked genes by the number of neighbors in the gene-level network. If a gene had no edge in the network, its rank was considered to be the number of genes with neighbors plus one. A gene’s rank-product is the product of its ranks from each network. The top general hub gene had the least rank-product.

To find hubs specific to a group of tissues, we used rank-product to rank hubs in both the target group of tissues, and all other tissues, separately. Then, we normalized ranks so that the top- and bottom-ranked hub have a score of 1 and 0, respectively. Let the normalized rank of a gene in the target group of tissues and other tissue be *r*_*t*_ and *r*_*o*_, respectively. Then, the F-score for the gene (*r*):

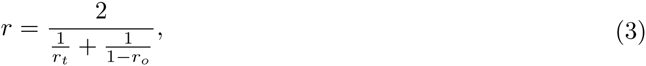

will be high if it ranks high in the target group, but low in other tissues.

We computed specific hubs for six groups of related tissues: 1) *skin– sun exposed* and *skin – not sun exposed*, 2) *adipose – subcutaneous*, *adipose – visceral* and *breast*, 3) *heart – left ventricle* and *skeletal muscle*, 4) *esophagus – mucosa* and *esophagus –muscularis*, 5) *artery – aorta* and *artery – tibial*, and 6) *nerve – tibial* and *artery – tibial*.

### TF-Target Enrichment in TE-TE edges

We downloaded transcription factors (TFs) and their known targets from ChEA (Lachmann et al., 2010). We measured the number of known TF-target relationships captured by a network, i.e., a TF and its target’s total expression nodes were directly connected with each other. We generated the null distribution of the number of known TF-target relationships by computing same test statistics for random networks, generated by permuting gene names among network nodes 1000 times. Then, we computed the empirical p-value as the proportion of those iterations for which the random network had at least as many known TF-Target edges as the test network. We fitted a Weibull distribution on the log(1+fraction of known TF-Target edges) to quantify the p-values.

### Per-pathway enrichment in TE-TE edges

We downloaded Reactome pathway genes from the Molecular Signature Database (c2.cp.reactome.v5.1) (Subramanian et al., 2005). For each Reactome pathway with at least ten genes, we tested whether the genes in the pathway had significantly smaller pairwise distances in our network than those in a random network, generated by permuting gene names among nodes in a network, using a Wilcoxon rank-sum test (Bonferroni corrected *p* ≤ 0.05).

### Same-pathway edge enrichment in TWN edges

We used Fisher’s exact test to check whether or not our networks were enriched with edges between genes from the same Reactome pathway. The null hypothesis was that two genes sharing an edge did not come from the same pathway.

### Tissue-specific networks (TSNs)

We built tissue-specific gene co-expression networks (TSNs) using an unsupervised Bayesian biclustering model, BicMix, on the gene level quantification from all of the GTEx v6 samples jointly (Gao et al., 2016). We performed 40 runs of BicMix on these data. We selected factors to build the tissue-specific covariance matrix estimate by including those for which non-zero factor values were exclusive to samples from the tissue of interest. We inverted these matrices and used GeneNet (Schafer and Strimmer, 2005), as in previous work, to build tissue-specific networks for each run (Gao et al., 2016). To build the final networks, we looked across each run with a network specific to a tissue, and every edge that appeared in > *r* runs was included in the final network. The parameter *r* was chosen for each tissue based on the number of runs with networks and the number of samples and genes in the run specific networks.

### Genotypes from GTEx data

The 2.5M and 5M BeadChip genotypes were merged to yield approximately 1.9 million genotyped SNPs. A greater set of genotypes were imputed using IMPUTE2 (Howie et al., 2009), yielding a satisfactory distribution of imputation scores for MAF ≥ 0.01 (mean INFO of 0.888 and median of 0.951 for variants with MAF between 0.01 and 0.05). The genotypes were filtered for MAF ≥ 0.05, leaving approximately 6 million variants. In order to take full advantage of SNP imputation, we used continuous (MLE of the dosage, ranging from 0 to 2) genotypes in association mapping. The genotype-level principle components were computed with the imputed genotypes.

### eQTL mapping

For each tissue, the gene expression values were projected to the quantiles of a standard normal, and Matrix-eQTL (Shabalin, 2012) was used to test all SNPs within the 150 Kb window of a gene’s transcription start site (TSS) or end site (TES) using an additive linear model. For cis-eQTL analyses, we optimized the number of PEER factors by tissue to a test chromosome (chromosome 11) to maximize the number of identified cis-eQTLs. We included in association mapping a tissue-specific number of PEER factors, sex, genotyping batch, and three genotype principal components. The correlation between SNP and gene expression levels was evaluated using the estimated t-statistic from this model. False discover rate (FDR) was calculated using the q-value R package (Storey, 2003; Dabney et al., 2010). We used these cis-eQTLs for the trans-eQTL analysis for the TSN edge replication.

### Trans-splicing QTLs

We computed trans-splicing QTLs using two approaches. In the first approach, we computed the best cis-associated variant per gene (smallest p-value) located within 1 Mb from the transcription start site (TSS) of the gene (Aguet et al., 2016). Then for every total expression node connected with an isoform ratio in the network, we measured association between the gene’s best cis-associated variant and all the isoform ratio neighbors using Matrix-eQTL (Shabalin, 2012), controlling for the first three genotype PCs and genotype platform. We corrected for false discovery (BH FDR ≤ 0.05). In the second approach, for each of the top 500 TE-IR hubs, we took all variants within 20 Kb of its TSS and tested their association with isoforms located in a different chromosome and connected with the total expression hub using Matrix-eQTL. Here, we used FDR ≤ 0.2 for multiple tests correction.

## Data Access

The Genotype-Tissue Expression v6 data are available through dbGaP, accession phs000424.v6 and the GTEx portal http://gtexportal.org. TWNs for 16 tissues, TSNs for 27 tissues, and four tissue subset networks are available for download at http://gtexportal.org (in progress).

Genotype, raw RNA-seq, quantified expression, and covariate data for the DGN cohort are available by application through the National Institute of Mental Health (NIMH) Center for Collaborative Genomic Studies on Mental Disorders. Instructions for requesting access to data can be found at the NIMH Repository and Genomics Resource (RGR; https://www.nimhgenetics.org/access_data_biomaterial.php), and inquiries should reference the “Depression Genes and Networks study (D. Levinson, PI)”.

## Acknowledgments

The Genotype-Tissue Expression (GTEx) Project was supported by the Common Fund of the Office of the Director of the National Institutes of Health. Additional funds were provided by the NCI, NHGRI, NHLBI, NIDA, NIMH, and NINDS. Donors were enrolled at Biospecimen Source Sites funded by NCI\SAIC-Frederick, Inc. (SAIC-F) subcontracts to the National Disease Research Interchange (10XS170), Roswell Park Cancer Institute (10XS171), and Science Care, Inc. (X10S172). The Laboratory, Data Analysis, and Coordinating Center (LDACC) was funded through a contract (HHSN268201000029C) to The Broad Institute, Inc. Biorepository operations were funded through an SAIC-F subcontract to Van Andel Institute (10ST1035). Additional data repository and project management were provided by SAIC-F (HHSN261200800001E). The Brain Bank was supported by a supplements to University of Miami grants DA006227 & DA033684 and to contract N01MH000028. Statistical Methods development grants were made to the University of Geneva (MH090941 & MH101814), the University of Chicago (MH090951, MH090937, MH101820, MH101825), the University of North Carolina-Chapel Hill (MH090936 & MH101819), Harvard University (MH090948), Stanford University (MH101782), Washington University St Louis (MH101810), and the University of Pennsylvania (MH101822). AB is supported by the Searle Scholars Program, NIH grant 1R01MH109905, NIH grant R01HG008150 (NHGRI; Non-Coding Variants Program), and NIH grant R01MH101814 (NIH Common Fund; GTEx Program). AG and BJ are funded by NIH grant 2T32HG003284-11. BEE is funded by NIH R00 HG006265, NIH R01 MH101822, NIH U01 HG007900, and a Sloan Faculty Fellowship.

## Disclosure Declaration

The authors declare no competing interests.

## References

Aguet, F., Brown, A. A., Castel, S., Davis, J. R., Mohammadi, P., Segre, A. V., Zappala, Z., Abell, N. S., Fresard, L., Gamazon, E. R., et al., 2016. Local genetic effects on gene expression across 44 human tissues. bioRxiv,.

Albert, R., 2005. Scale-free networks in cell biology. Journal of Cellular Science, 118(Pt 21):4947–4957.

Alexa, A. and Rahnenfuhrer, J., 2016. topGO: Enrichment Analysis for Gene Ontology.

Andrews, S., 2010. FastQC: a quality control tool for high throughput sequence data.

Ashburner, M., Ball, C. A., Blake, J. A., Botstein, D., Butler, H., Cherry, J. M., Davis, A. P., Dolinski, K., Dwight, S. S., Eppig, J. T., et al., 2000. Gene Ontology: tool for the unification of biology. Nature Genetics, 25(1):25–29.

Auboeuf, D., Dowhan, D. H., Li, X., Larkin, K., Ko, L., Berget, S. M., and O’Malley, B. W., 2004. CoAA, a Nuclear Receptor Coactivator Protein at the Interface of Transcriptional Coactivation and RNA Splicing. Molecular and Cellular Biology, 24(1):442–453.

Auboeuf, D., Hönig, A., Berget, S. M., and O’Malley, B. W., 2002. Coordinate regulation of transcription and splicing by steroid receptor coregulators. Science, 298(5592):416–9.

Ballouz, S., Verleyen, W., and Gillis, J., 2015. Guidance for RNA-seq co-expression network construction and analysis: Safety in numbers. Bioinformatics, 31(13):2123–2130.

Barabasi, A. L. and Oltvai, Z. N., 2004. Network biology: Understanding the cell’s functional organization [Review]. Nature Reviews Genetics, 5(2):101–NIL.

Battle, A., Mostafavi, S., Zhu, X., Potash, J. B., Weissman, M. M., McCormick, C., Haudenschild, C. D., Beckman, K. B., Shi, J., Mei, R., et al., 2014. Characterizing the genetic basis of transcriptome diversity through RNA-sequencing of 922 individuals. Genome Research, 24(1):14–24.

Blencowe, B. J., Baurén, G., Eldridge, A. G., Issner, R., Nickerson, J. A., Rosonina, E., and Sharp, P. A., 2000. The SRm160/300 splicing coactivator subunits. RNA, 6(1):111–20.

Bolger, A. M., Lohse, M., and Usadel, B., 2014. Trimmomatic: A flexible trimmer for Illumina sequence data. Bioinformatics,:btu170.

Buettner, F., Natarajan, K. N., Casale, F. P., Proserpio, V., Scialdone, A., Theis, F. J., Teichmann, S. A., Marioni, J. C., and Stegle, O., 2015. Computational analysis of cell-to-cell heterogeneity in single-cell RNA-sequencing data reveals hidden subpopulations of cells. Nature Biotechnology, 33(2):155–160.

Chen, M. and Manley, J. L., 2009. Mechanisms of alternative splicing regulation: insights from molecular and genomics approaches. Nature Reviews Molecular Cell Biology, 10(11):741–54.

Dabney, A., Storey, J. D., and Warnes, G., 2010. qvalue: Q-value estimation for false discovery rate control. R Package Version, 1(0).

Dai, C., Li, W., Liu, J., and Zhou, X. J., 2012. Integrating many co-splicing networks to reconstruct splicing regulatory modules. BMC Systems Biology, 6(Suppl 1):S17.

DeBoever, C., Ghia, E. M., Shepard, P. J., Rassenti, L., Barrett, C. L., Jepsen, K., Jamieson, C. H., Carson, D., Kipps, T. J., and Frazer, K. A., et al., 2015. Transcriptome sequencing reveals potential mechanism of cryptic 3’ splice site selection in *SF3B1* -mutated cancers. PLoS Computational Biology, 11(3):e1004105.

Derrien, T., Estellé, J., Sola, S. M., Knowles, D. G., Raineri, E., Guigó, R., and Ribeca, P., 2012. Fast computation and applications of genome mappability. PLoS ONE, 7(1).

Dobin, A., Davis, C. A., Schlesinger, F., Drenkow, J., Zaleski, C., Jha, S., Batut, P., Chaisson, M., and Gingeras, T. R., 2013. STAR: Ultrafast universal RNA-seq aligner. Bioinformatics, 29(1):15–21.

Du, C., Ma, X., Meruvu, S., Hugendubler, L., and Mueller, E., 2014. The adipogenic transcriptional cofactor ZNF638 interacts with splicing regulators and influences alternative splicing. Journal of Lipid Research, 55(9):1886–96.

D’Souza, I., Poorkaj, P., Hong, M., Nochlin, D., Lee, V. M.-Y., Bird, T. D., and Schellenberg, G. D., 1999. Missense and silent tau gene mutations cause frontotemporal dementia with parkinsonism-chromosome 17 type, by affecting multiple alternative RNA splicing regulatory elements. Proceedings of the National Academy of Sciences, 96(10):5598–5603.

Fabregat, A., Sidiropoulos, K., Garapati, P., Gillespie, M., Hausmann, K., Haw, R., Jassal, B., Jupe, S., Korninger, F., McKay, S., et al., 2016. The Reactome pathway knowledgebase. Nucleic Acids Research, 44(D1):D481–D487.

Friedman, J., Hastie, T., and Tibshirani, R., 2008. Sparse inverse covariance estimation with the graphical lasso. Biostatistics, 9(3):432–441.

Gao, C., McDowell, I. C., Zhao, S., Brown, C. D., and Engelhardt, B. E., 2016. Context specific and differential gene co-expression networks via Bayesian biclustering. PLoS Computational Biology, 12(7):e1004791.

Gao, G., Dudley, S. C., and Jr., 2013. RBM25/LUC7L3 function in cardiac sodium channel splicing regulation of human heart failure. Trends in Cardiovascular Medicine, 23(1):5–8.

Ghigna, C., Valacca, C., and Biamonti, G., 2008. Alternative splicing and tumor progression. Current Genomics, 9(8):556–570.

Glatz, D. C., Rujescu, D., Tang, Y., Berendt, F. J., Hartmann, A. M., Faltraco, F., Rosenberg, C., Hulette, C., Jellinger, K., Hampel, H., et al., 2006. The alternative splicing of tau exon 10 and its regulatory proteins CLK2 and TRA2-BETA1 changes in sporadic Alzheimer’s disease. Journal of Neurochemistry, 96(3):635–644.

Greene, C. S., Krishnan, A., Wong, A. K., Ricciotti, E., Zelaya, R. A., Himmelstein, D. S., Zhang, R., Hartmann, B. M., Zaslavsky, E., Sealfon, S. C., et al., 2015. Understanding multicellular function and disease with human tissue-specific networks. Nature Genetics, 47(6):569–576.

GTEx Consortium, 2015. The Genotype-Tissue Expression (GTEx) pilot analysis: Multitissue gene regulation in humans. Science, 348(6235):648–660.

Hormozdiari, F., Penn, O., Borenstein, E., and Eichler, E. E., 2015. The discovery of integrated gene networks for autism and related disorders. Genome Research, 25:142–154.

Howie, B. N., Donnelly, P., and Marchini, J., 2009. A flexible and accurate genotype imputation method for the next generation of genome-wide association studies. PLoS Genetics, 5(6).

Hsieh, C.-J., Sustik, M. A., Dhillon, I. S., and Ravikumar, P., 2011. Sparse Inverse Covariance Matrix Estimation Using Quadratic Approximation. Advances in Neural Information Processing Systems, 24:2330–2338.

Hutton, M., Lendon, C. L., Rizzu, P., Baker, M., Froelich, S., Houlden, H., Pickering-Brown, S., Chakraverty, S., Isaacs, A., Grover, A., et al., 1998. Association of missense and 5’-splice-site mutations in tau with the inherited dementia FTDP-17. Nature, 393(6686):702–705.

Iancu, O. D., Colville, A., Oberbeck, D., Darakjian, P., McWeeney, S. K., and Hitzemann, R., 2015. Cosplicing network analysis of mammalian brain RNA-Seq data utilizing WGCNA and Mantel correlations. Frontiers in Genetics, 6.

Jeong, H., Mason, S. P., Barabási, a. L., and Oltvai, Z. N., 2001. Lethality and centrality in protein networks. Nature, 411(6833):41–42.

Jo, B., He, Y., Strober, B. J., Parsana, P., Aguet, F., Brown, A. A., Castel, S. E., Gamazon, E. R., Gewirtz, A., Gliner, G., et al., 2016. Distant regulatory effects of genetic variation in multiple human tissues. bioRxiv,:074419.

Khatri, P., Sirota, M., and Butte, A. J., 2012. Ten years of pathway analysis: Current approaches and outstanding challenges. PLoS Computational Biology, 8(2).

Kulisz, A. and Simon, H.-G., 2008. An evolutionarily conserved nuclear export signal facilitates cytoplasmic localization of the Tbx5 transcription factor. Molecular and Cellular Biology, 28(5):1553–64.

Kumar P., P., Franklin, S., Emechebe, U., Hu, H., Moore, B., Lehman, C., Yandell, M., Moon, A. M., Rodriguez, M., Aladowicz, E., et al., 2014. TBX3 Regulates Splicing In Vivo: A Novel Molecular Mechanism for Ulnar-Mammary Syndrome. PLoS Genetics, 10(3):e1004247.

Lachmann, A., Xu, H., Krishnan, J., Berger, S. I., Mazloom, A. R., and Ma’ayan, A., 2010. ChEA: transcription factor regulation inferred from integrating genome-wide ChIP-X experiments. Bioinformatics, 26(19):2438–2444.

Langmead, B., Trapnell, C., Pop, M., and Salzberg, S., 2009. 2C-Ultrafast and memory-efficient alignment of short DNA sequences to the human genome. Genome Biology, 10(3):R25.

Lee, H. K., Hsu, A. K., Sajdak, J., Qin, J., and Pavlidis, P., 2004. Coexpression analysis of human genes across many microarray data sets. Genome Research, 14(6):1085–1094.

Lee, Y., Gamazon, E. R., Rebman, E., Lee, Y., Lee, S., Dolan, M. E., Cox, N. J., and Lussier, Y. A., 2012. Variants Affecting Exon Skipping Contribute to Complex Traits. PLoS Genetics, 8(10).

Leek, J. T., Scharpf, R. B., Bravo, H. C., Simcha, D., Langmead, B., Johnson, W. E., Geman, D., Baggerly, K., and Irizarry, R. A., 2010. Tackling the widespread and critical impact of batch effects in high-throughput data. Nature Reviews Genetics, 11(10):733–739.

Li, B. and Dewey, C. N., 2011. RSEM: accurate transcript quantification from RNA-Seq data with or without a reference genome. BMC Bioinformatics, 12(1):1.

Li, C., Lin, R.-I., Lai, M.-C., Ouyang, P., and Tarn, W.-Y., 2003. Nuclear Pnn/DRS protein binds to spliced mRNPs and participates in mRNA processing and export via interaction with RNPS1. Molecular and Cellular Biology, 23(20):7363–76.

Li, H.-D., Menon, R., Eksi, R., Guerler, A., Zhang, Y., Omenn, G. S., and Guan, Y., 2016a. A Network of Splice Isoforms for the Mouse. Scientific Reports, 6:24507.

Li, H.-D., Omenn, G. S., and Guan, Y., 2015. MIsoMine: a genome-scale high-resolution data portal of expression, function and networks at the splice isoform level in the mouse. Database: The Journal of Biological Databases and Curation, 2015:bav045.

Li, W., Kang, S., Liu, C.-C., Zhang, S., Shi, Y., Liu, Y., and Zhou, X. J., 2014. High-resolution functional annotation of human transcriptome: predicting isoform functions by a novel multiple instance-based label propagation method. Nucleic Acids Research, 42(6):e39.

Li, Y. I., van de Geijn, B., Raj, A., Knowles, D. A., Petti, A. A., Golan, D., Gilad, Y., and Pritchard, J. K., 2016b. Rna splicing is a primary link between genetic variation and disease. Science, 352(6285):600–604.

Liu, Q., Gao, J., Chen, X., Chen, Y., Chen, J., Wang, S., Liu, J., Liu, X., and Li, J., 2008. HBP21: a novel member of TPR motif family, as a potential chaperone of heat shock protein 70 in proliferative vitreoretinopathy (PVR) and breast cancer. Molecular Biotechnology, 40(3):231–240.

López-Bigas, N., Audit, B., Ouzounis, C., Parra, G., and Guigó, R., 2005. Are splicing mutations the most frequent cause of hereditary disease? FEBS Letters, 579(9):1900–1903.

Magomedova, L., Tiefenbach, J., Zilberman, E., Voisin, V., Robitaille, M., Gueroussov, S., Irimia, M., Ray, D., Patel, R., Xu, C., et al., 2016. ARGLU1 is a Glucocorticoid Receptor Coactivator and Splicing Modulator Important in Stress Hormone Signaling and Brain Development. bioRxiv,.

Martínez-Redondo, V., Jannig, P. R., Correia, J. C., Ferreira, D. M. S., Cervenka, I., Lindvall, J. M., Sinha, I., Izadi, M., Pettersson-Klein, A. T., Agudelo, L. Z., et al., 2016. Peroxisome proliferator-activated receptor gamma coactivator-1 alpha isoforms selectively regulate multiple splicing events on target genes. Journal of Biological Chemistry, 291(29):15169–15184.

Matlin, A. J., Clark, F., and Smith, C. W. J., 2005. Understanding alternative splicing: towards a cellular code. Nature Reviews Molecular Cell Biology, 6(5):386–98.

McDowell, I. C., Pai, A. A., Guo, C., Vockley, C. M., Brown, C. D., Reddy, T. E., and Engelhardt, B. E., 2016. Many long intergenic non-coding RNAs distally regulate mRNA gene expression levels. bioRXiv preprint 044719,.

Melé, M., Ferreira, P. G., Reverter, F., DeLuca, D. S., Monlong, J., Sammeth, M., Young, T. R., Goldmann, J. M., Pervouchine, D. D., Sullivan, T. J., et al., 2015. The human transcriptome across tissues and individuals. Science, 348(6235):660–665.

Mostafavi, S., Battle, A., Zhu, X., Potash, J., Weissman, M., Shi, J., Beckman, K., Haudenschild, C., McCormick, C., Mei, R., et al., 2014. Type I interferon signaling genes in recurrent major depression: increased expression detected by whole-blood RNA sequencing. Molecular Psychiatry, 19(12):1267–1274.

Mostafavi, S., Battle, A., Zhu, X., Urban, A. E., Levinson, D., Montgomery, S. B., and Koller, D., 2013. Normalizing RNA-sequencing data by modeling hidden covariates with prior knowledge. PloS ONE, 8(7):e68141.

Mullick, A., Leon, Z., Min-Oo, G., Berghout, J., Lo, R., Daniels, E., and Gros, P., 2006. Cardiac Failure in C5-Deficient A/J Mice after Candida albicans Infection. Infection and Immunity, 74(8):4439–4451.

Ong, C.-T. and Corces, V. G., 2011. Enhancer function: new insights into the regulation of tissue-specific gene expression. Nature Reviews Genetics, 12(4):283–93.

Penrod, N. M., Cowper-Sal-Lari, R., and Moore, J. H., 2011. Systems genetics for drug target discovery. Trends in Pharmacological Sciences, 32(10):623–630.

Pierson, E., Koller, D., Battle, A., Mostafavi, S., Consortium, G., et al., 2015. Sharing and specificity of co-expression networks across 35 human tissues. PLoS Computational Biology, 11(5):e1004220.

Piro, R. M., Ala, U., Molineris, I., Grassi, E., Bracco, C., Perego, G. P., Provero, P., and Di Cunto, F., 2011. An atlas of tissue-specific conserved coexpression for functional annotation and disease gene prediction. European Journal of Human Genetics, 19(11):1173–1180.

Prieto, C., Risueño, A., Fontanillo, C., and De Las Rivas, J., 2008. Human gene coexpression landscape: Confident network derived from tissue transcriptomic profiles. PLoS ONE, 3(12).

Qian, J., Esumi, N., Chen, Y., Wang, Q., Chowers, I., and Zack, D. J., 2005. Identification of regulatory targets of tissue-specific transcription factors: application to retina-specific gene regulation. Nucleic Acids Research, 33(11):3479–3491.

Roider, H. G., Manke, T., O’keeffe, S., Vingron, M., and Haas, S. A., 2009. PASTAA: Identifying transcription factors associated with sets of co-regulated genes. Bioinformatics, 25(4):435–442.

Rue, H. and Held, L., 2005. Gaussian Markov Random Fields: Theory and Applications. Monographs on Statistics and Applied Probability. Chapman & Hall, London.

Saito, Y., Kojima, T., and Takahashi, N., 2012. Mab21l2 is essential for embryonic heart and liver development. PLOS One, 7(3):e32991.

Schäfer, J. and Strimmer, K., 2005. A shrinkage approach to large-scale covariance matrix estimation and implications for functional genomics. Statistical Applications in Genetics and Molecular Biology, 4:Article32.

Schafer, J. and Strimmer, K., 2005. An empirical Bayes approach to inferring large-scale gene association networks. Bioinformatics, 21(6):754–764.

Scotti, M. M. and Swanson, M. S., 2015. RNA mis-splicing in disease. Nature Reviews Genetics, 17(1):19–32.

Shabalin, A. A., 2012. Matrix eQTL: ultra fast eQTL analysis via large matrix operations. Bioinformatics, 28(10):1353–8.

Silver, D. L., Watkins-Chow, D. E., Schreck, K. C., Pierfelice, T. J., Larson, D. M., Burnetti, A. J., Liaw, H.-J., Myung, K., Walsh, C. A., Gaiano, N., et al., 2010. The exon junction complex component Magoh controls brain size by regulating neural stem cell division. Nature Neuroscience, 13(5):551–558.

Squier, C. A. and Kremer, M. J., 2001. Biology of oral mucosa and esophagus. Journal of the National Cancer Institute. Monographs, 2001(29):7–15.

Storey, J. D., 2003. The positive false discovery rate: A Bayesian interpretation and the q-value. Annals of Statistics,:2013–2035.

Stuart, J. M., Segal, E., Koller, D., and Kim, S. K., 2003. R ESEARCH A RTICLES A Gene-Coexpression Network. Science, 302(5643):249–255.

Subramanian, A., Tamayo, P., Mootha, V. K., Mukherjee, S., Ebert, B. L., Gillette, M. A., Paulovich, A., Pomeroy, S. L., Golub, T. R., Lander, E. S., et al., 2005. Gene set enrichment analysis: A knowledge-based approach for interpreting genome-wide expression profiles. Proceedings of the National Academy of Sciences, 102(43):15545–15550.

Sui, Y., Yang, Z., Xiong, S., Zhang, L., Blanchard, K. L., Peiper, S. C., Dynan, W. S., Tuan, D., and Ko, L., 2007. Gene amplification and associated loss of 5’ regulatory sequences of CoAA in human cancers. Oncogene, 26(6):822–835.

Sveen, A., Kilpinen, S., Ruusulehto, A., Lothe, R., and Skotheim, R., 2015. Aberrant RNA splicing in cancer; expression changes and driver mutations of splicing factor genes. Oncogene,.

Tomsic, J., He, H., Akagi, K., Liyanarachchi, S., Pan, Q., Bertani, B., Nagy, R., Symer, D. E., Blencowe, B. J., and de la Chapelle, A., et al., 2015. A germline mutation in SRRM2, a splicing factor gene, is implicated in papillary thyroid carcinoma predisposition. Scientific Reports, 5:10566.

Vareli, K., Frangou-Lazaridis, M., van der Kraan, I., Tsolas, O., and van Driel, R., 2000. Nuclear distribution of prothymosin alpha and parathymosin: evidence that prothymosin alpha is associated with RNA synthesis processing and parathymosin with early DNA replication. Experimental Cell Research, 257(1):152–61.

Wang, E. T., Sandberg, R., Luo, S., Khrebtukova, I., Zhang, L., Mayr, C., Kingsmore, S. F., Schroth, G. P., and Burge, C. B., 2008. Alternative isoform regulation in human tissue transcriptomes. Nature, 456(7221):470–6.

Wang, Z. and Burge, C. B., 2008. Splicing regulation: from a parts list of regulatory elements to an integrated splicing code. RNA, 14(5):802–13.

Ward, A. J. and Cooper, T. A., 2010. The pathobiology of splicing. The Journal of Pathology, 220(2):152–163.

Warde-Farley, D., Donaldson, S. L., Comes, O., Zuberi, K., Badrawi, R., Chao, P., Franz, M., Grouios, C., Kazi, F., Lopes, C. T., et al., 2010. The GeneMANIA prediction server: biological network integration for gene prioritization and predicting gene function. Nucleic Acids Research, 38(suppl 2):W214–W220.

Weiser, M., Mukherjee, S., and Furey, T. S., 2014. Novel distal eqtl analysis demonstrates effect of population genetic architecture on detecting and interpreting associations. Genetics, 198(3):879–893.

Westra, H.-J., Peters, M. J., Esko, T., Yaghootkar, H., Schurmann, C., Kettunen, J., Christiansen, M. W., Fairfax, B. P., Schramm, K., Powell, J. E., et al., 2013. Systematic identification of trans eQTLs as putative drivers of known disease associations. Nature Genetics, 45(10):1238–43.

Witten, J. T. and Ule, J., 2011. Understanding splicing regulation through RNA splicing maps. Trends in Genetics, 27(3):89–97.

Wu, J. Y., Kar, A., Kuo, D., Yu, B., and Havlioglu, N., 2006. SRp54 (SFRS11), a regulator for tau exon 10 alternative splicing identified by an expression cloning strategy. Molecular and Cellular Biology, 26(18):6739–47.

Xiao, X., Moreno-moral, A., Rotival, M., Bottolo, L., and Petretto, E., 2014. Multi-tissue Analysis of Co-expression Networks by Higher-Order Generalized Singular Value Decomposition Identifies Functionally Coherent Transcriptional Modules. PLoS Genetics, 10(1).

Yang, Y., Han, L., Yuan, Y., Li, J., Hei, N., and Liang, H., 2014. Gene co-expression network analysis reveals common system-level properties of prognostic genes across cancer types. Nature Communications, 5:1–9.

Zhang, B. and Horvath, S., 2005. A general framework for weighted gene co-expression network analysis. Statistical Applications in Genetics and Molecular Biology, 4:Article17.

Zhang, W. J. and Wu, J. Y., 1996. Functional properties of p54, a novel SR protein active in constitutive and alternative splicing. Molecular and Cellular Biology, 16(10):5400–8.

Zhong, R., Allen, J. D., Xiao, G., and Xie, Y., 2014. Ensemble-based network aggregation improves the accuracy of gene network reconstruction. PLoS ONE, 9(11):1–10.

Zhou, A., Ou, A. C., Cho, A., Benz, E. J., and Huang, S.-C., 2008. Novel splicing factor RBM25 modulates Bcl-x pre-mRNA 5’ splice site selection. Molecular and Cellular Biology, 28(19):5924–36.

Zimowska, G., Shi, J., Munguba, G., Jackson, M. R., Alpatov, R., Simmons, M. N., Shi, Y., and Sugrue, S. P., 2003. Pinin/DRS/memA Interacts with SRp75, SRm300 and SRrp130 in Corneal Epithelial Cells. Investigative Opthalmology & Visual Science, 44(11):4715.

